# *spiralize*: an R Package for Visualizing Data on Spirals

**DOI:** 10.1101/2021.06.30.450482

**Authors:** Zuguang Gu, Daniel Hübschmann

## Abstract

**Summary:** Spiral layout has two major advantages for data visualization. First, it is able to visualize data with long axes, which greatly improves the resolution of visualization. Second, it is efficient for time series data to reveal periodic patterns. Here we present the R package *spiralize* that provides a general solution for visualizing data on spirals. *spiralize* implements numerous graphics functions so that self-defined high-level graphics can be easily implemented by users. The flexibility and power of *spiralize* are demonstrated by five examples from real-world datasets.

**Availability and implementation:** The *spiralize* package and documentations are freely available at the Comprehensive R Archive Network (CRAN) https://CRAN.R-project.org/package=spiralize.

**Supplementary information:** Supplementary data are available at *Bioinformatics* online.

## 1 Introduction

High resolution visualization is always an urgent need for data analysis in general and biological data in particular. In a figure with a width of 1000 pixels, traditional linear visualization can only maximally distinguish 1000 data points on the *x*-axis, which restricts visualization of larger datasets. There are methods aiming to improve the resolution of visualization by extending the *x*-axis into two-dimensional space by certain transformations. Circular visualization (Gu *et al*., 2014) extends the axis to a circle, which improves the resolution by a small factor of 3.14 and allows multiple tracks for simultaneous visualization of different data types. Hilbert curve (Gu *et al*., 2016) is another type of method that densely folds the axis into two-dimensional space and it greatly improves the resolution from *n* to *n*^2^. However, it is only efficient for visualizing a single data type on the (one-dimensional) folded axis. Visualization on spirals is an alternative method (we refer to Archimedean spirals in this work) where the *x*-axis is mapped to a spiral. With a large number of loops, the spiral will have a large length to allow more data points, thus it can visualize data with high resolution. On the other hand, if there are fewer loops, although less data points can be put on the spiral, there are more spaces between loops, thus it is possible to perform multi-track visualization along the spiral. The second advantage of a spiral is its efficiency to visualize time series datasets to reveal periodic patterns because of its cyclic characteristic. Indeed, in many current studies, spirals were mainly used for visualizing time series datasets (Weber *et al*., 2001; Carlis and Konstan, 1998). An alternative application was to visualize trees of life of more than 50000 species (Hedges *et al*., 2015).

In polar coordinates (*r, θ*), an Archimedean spiral has the form *r* = *bθ* with *b* as the free parameter. The radial distance between two neighbouring loops is calculated as *d* = 2*πb* for all *θ*. This implies the radial distance is always constant regardless of the positions on the spiral, thus it is very suitable to put tracks on spirals where the radial direction corresponds to the *y*-axis. Spiral visualization has higher resolution than traditional linear visualization. For example, a spiral with 4 loops improves the resolution of visualization by a factor of 6.3 and a spiral with 10 loops improves by a factor of 15.7. In Supplementary File 1, we demonstrated the improvement of resolution increases almost linearly to the number of loops in the spiral. However, users need to be cautious when selecting more loops in spiral visualization as there will be less space between two neighbouring loops, thus less graphics can be conveniently arranged in the spiral.

In this work, we developed a new R package *spiralize*, which provides a general solution for data visualization on spirals. *spiralize* provides numerous low-level graphics functions so that user-defined complex plots can be easily constructed by combining them. We demonstrate the use of *spiralize* by five examples based on real-world datasets and we believe *spiralize* will greatly facilitate better data.

## 2 Implementation

*spiralize* supports multiple tracks along the spiral where each track serves as an independent plotting region. In each track, numerous low-level graphics functions implemented in *spiralize* can be used to add basic graphics, *e.g*., points, lines, polygons, texts, axes and images, to build complex plots. *spiralize* also provides functions to draw dendrograms or phylogenetic trees with huge numbers of leaves on the spiral.

Horizon chart is a visualization method that vertically splits an area chart with uniform size, then the bands are layered on top of each other (Heer *et al*., 2009). It reduces the height of an area chart by a factor of *k* times where *k* is the number of bands, thus it is very suitable for spiral visualization because tracks normally have small heights on spirals. Furthermore, it has already been applied in a previous study that utilized spiral visualization (Tominski and Schumann, 2008). *spiralize* specifically implements a function `spiral_horizon()` to draw horizon charts along the spiral; its usage is demonstrated in Figure 1A and 1C.

**Figure 1:**
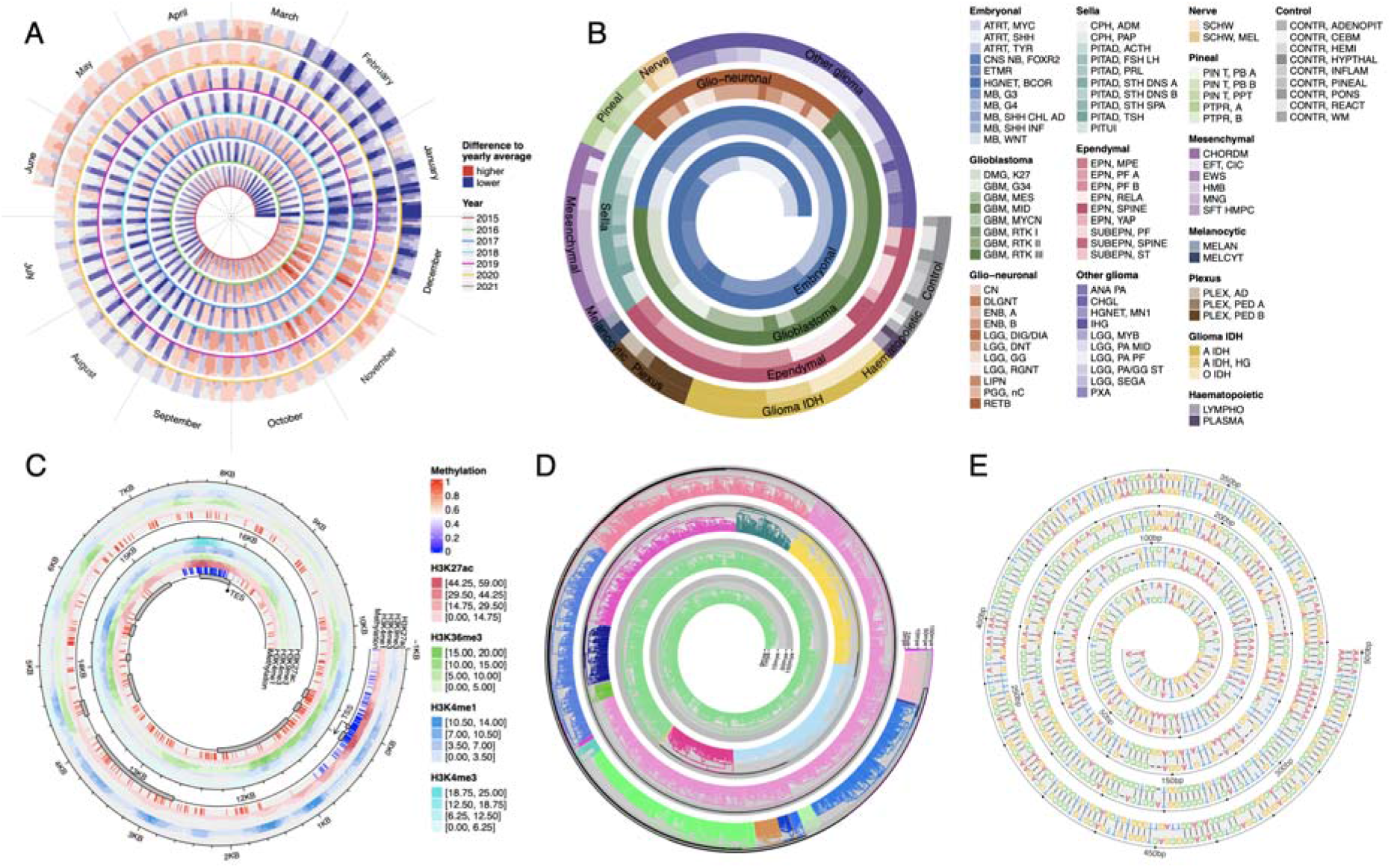
Examples of *spiralize* visualizations. A) Daily downloads of the *ggplot2* package. B) 91 subtypes of central nervous system tumors from 2801 samples. C) DNA Methylation and four histone modification signals along the gene *BCAT2*. D) Phylogenetic tree of 3738 mammals. Unit on the *y*-axis: million years ago (mya). E) Pairwise alignment of coding sequence of the gene *TP53* between human and its mouse homologue. Source code for generating all figures can be found in Supplementary File 2. *spiralize* provides more real-world examples in the vignettes.

*spiralize* supports two methods for mapping data to spirals for different scenarios of data analysis. The method “angle” linearly maps data to angles in the polar coordinate system, whereas the method “curve” linearly maps data to the length of the spiral curve measured at corresponding points. With the “angle” method, it is possible to directly compare between loops to establish corresponding patterns at the same phase across different periods (*e.g*., Figure 1A), thus it is more suitable for time series datasets. However, the drawback is that intervals with equal bin sizes of data are visually wider in outer loops than in inner loops. As a comparison, the “curve” method does not illustrate any periodic pattern but it keeps the interval widths consistent if they correspond to equal bin sizes of the data. (*e.g*., Figure 1B-1E).

## 3 Application

Figure 1A visualizes daily downloads of the R package *ggplot2* from 2015-01-01 to 2021-06-14. Values on the plot are log2-values of the ratio between daily downloads and average downloads of the corresponding year. In Figure 1A, the “angle” method was used to map data and each loop of the spiral corresponds to 52 weeks. In this way, the same calendar weeks of different years can be correctly aligned and compared. Figure 1A clearly illustrates that there are fewer downloads during weekends and vacation times, *e.g*., end of December, beginning of January and summer vacations (*i.e*., July and August). For time series data, *spiralize* automatically calculates the start and end angles on the spiral to make the positions of the two corresponding time points in the loop reflect its proportion in the corresponding period (Note the position of the end angle in Figure 1A).

Figure 1B visualizes a subtype classification of central nervous system tumors (Capper *et al*., 2018). The dataset contains 14 different tumor types that are classified into 91 subtypes based on DNA methylation from 2801 samples. On the spiral, all subtypes can be easily distinguished and the relative fractions of subtypes in the cohort can be visually identified by the widths of corresponding bars. These details are difficult to reveal by traditional visualization methods.

Figure 1C visualizes DNA methylation and signals of four histone modifications along the gene *BCAT2* in human lung tissue, which covers a genomic range of around 18kb where *BCAT2* is extended by 1kb both 5’ and 3’. The data was obtained from the GEO database with accession ID GSE16256 (Bernstein *et al*., 2010). The plot clearly illustrates the correspondence between various epigenomics signals. The resolution can reach almost 10 base pairs per pixel in a figure with size 7×7 inches.

Figure 1D visualizes a phylogenetic tree of 3738 mammal species (Hedges *et al*., 2015). The tree was cut into 26 branches and each branch was assigned a different color. On the *y*-axis, the heights of nodes in the tree which correspond to the evolutionary distances were log10-transformed. In the vignette of the *spiralize* package, we reimplemented the complete phylogenetic tree from Hedges *et al*., 2015 which contains 50455 species.

Figure 1E visualizes pairwise alignment of coding sequences (CDS) of the gene *TP53* between human and its mouse homologue. The alignment was obtained by blast. Due to the space of the plot in Figure 1E, only the first 500 base pairs are shown. The full set of 1194 base pairs of the alignment can be found in the vignette of the *spiralize* package.

