## Supplementary File 1 for "*spiralize*: an R Package for Visualizing Data on Spirals": supplementary1.html (from another version of this preprint: version 1, 2021-07-01)

Supplementary file 1. Resolution of the spiral plot


### Supplementary file 1. Resolution of the spiral plot

###### Zuguang Gu

#### 2021-06-30

In polar coordinates (\(r\), \(\theta\)), the Archimedean spiral has the following form:

\[ r = b \cdot \theta \]

The length of the spiral denoted as \(l\) is calculated as:

\[ l = \frac{b \cdot \mathrm{ln}\left(\theta + \sqrt{1 + \theta^2}\right) + b \cdot \theta\sqrt{1 + \theta^2}}{2} \]

In the plot above, denote the maximal radius of the spiral as \(d\_{max} = b \cdot \theta\_{max}\). We can treat \(2d\_{max}\) as the resolution of the visualization applied in the normal Cartesian coordinate system and \(l\_{max} = l(\theta\_{max})\) as the resolution of the visualization applied on the spiral. Then the ratio of the two resolutions is:

\[ ratio = \frac{l\_{max}}{2 \cdot d\_{max}}
= \frac{\mathrm{ln}\left(\theta\_{max} + \sqrt{1 + \theta\_{max}^2}\right) + \theta\_{max}\sqrt{1 + \theta\_{max}^2}}{4\theta\_{max}} \]

E.g., for a spiral with 4 loops (\(\theta\_{max} = 8\pi\)), the ratio is 6.33, which means the spiral improves the resolution of visualization by a factor of more than 6 times. Generally, the ratio increases almost linearly to the number of loops in the spiral.
